# Decoding Numeracy and Literacy in the Human Brain: Insights from MEG and MVPA

**DOI:** 10.1101/2022.12.29.522254

**Authors:** Sanjeev Nara, Haider Raza, Manuel Carreiras, Nicola Molinaro

## Abstract

Numbers and letters are the fundamental building blocks of our everyday social interactions. Previous studies have focused on determining the cortical networks shaped by numeracy and literacy in the human brain, partially supporting the hypothesis of distinct neural circuits involved the processing of the two categories. In this study, we aim to investigate the temporal dynamics for number and letter processing. We present MEG data from two experiments (N=20 each). In the first experiment, single numbers, letter, and their respective false fonts (false numbers and false letters) were presented, whereas, in the second experiment, numbers, letters, and their respective false fonts were presented as a string of characters. We used multivariate pattern analysis techniques (time-resolved decoding and temporal generalization) testing the strong hypothesis that the neural mechanisms supporting letter and number processing can be logistically classified as categorically separate. Our results show a very early dissociation (∼100 ms) between numbers, and letters when compared to false fonts. Number processing can be dissociated with similar accuracy when presented as isolated items or strings of characters, while letter processing shows dissociable classification accuracy for single items compared to strings. These findings reinforce the evidence indicating that early visual processing can be differently shaped by the experience with numbers and letters; this dissociation is stronger for strings compared to single items, thus showing that combinatorial mechanisms for numbers and letters could be categorically distinguished and influence early brain activity.

## Introduction

Numbers and letters are fundamental building blocks of our everyday social interactions. Humans constantly mix alphanumeric codes in their everyday language experience. Such skills depend on the acquisition of literacy and numeracy, which are fundamental steps in standard education. They are formally acquired in the first years of primary school, so experience with alphanumeric codes develops in a similar developmental window. While literacy acquisition changes the human brain circuitry [1]–[3], evidence on numeracy stimulating an independent brain network is still debated. In the present study, we evaluate if the human brain responses to alphabetic and numerical symbols can be categorically classified, providing thus strong evidence for the hypothesis that literacy and numeracy shape distinct neuronal circuits.

A large number of studies focused on the definition of the brain networks and the neural biomarkers, related to numbers and letter processing. Using iEEG (intracranial electroencephalography), Shum and colleagues (2013) [4] isolated an inferior temporal region in the right hemisphere that responded preferentially to numbers. In an fMRI study, Park and colleagues (2012) reported opposite hemispheric recruitment for numbers and letters [5]: strings of numbers recruited mainly the right lateral occipital regions, while strings of consonants recruited more the left mid-fusiform and the inferior temporal regions. In a follow-up ERP study, Park and colleagues (2014) [6] observed a similar division of labor for the two hemispheres concerning the location of the N100 component (mainly between 140 and 170 msec), more evident in left-lateralized electrodes for letters and more right-lateralized for numbers. Despite it is not possible to determine an unambiguous brain source for these ERP effects, this is evidence for the recruitment of different brain networks for the two types of visual stimuli. The string of letters elicited larger P200 responses “*suggesting that the visual cortex is tuned to selectively process combinations of letters, but not numbers, further along in the visual processing stream*”. In this last study, the authors also explored the neural response to single stimuli compared to strings of stimuli and no reliable difference was observed for single numbers compared to single letters. Importantly, such strong experiential influence of reading and mathematics on the human visual system would mature during primary school and would be later evident in adolescence and adulthood [7].

A different line of studies however, brought evidence supporting the idea that numerals recruit populations of neurons in both hemispheres, bilaterally, providing evidence against the unique right hemispheric specialization. For example, in a MEG study, Carreiras and colleagues (2015) [8] reported bilateral recruitment for strings of numbers at around 150 ms compared to other alphabetic string conditions (consonant strings, words, and pseudowords), which mainly showed activation in the left hemisphere. In an fMRI study, Grotheer and colleagues (2016) [9] observed a bilateral preferential response for numbers in the inferior temporal gyrus (ITG) when compared with letters, false numbers, or everyday objects. These studies suggest that rather than one specific hemisphere or brain area, numeracy and literacy are complex processes and recruit an extended network of highly overlapping brain regions, whose lateralization would not be so clear-cut. In other words, similar regions (especially in the left hemisphere) could be recruited quantitatively more or less for numbers or letters, and the related neural activity could not be easily dissociated. To address this issue, here we focused on a multivariate pattern analysis (MVPA) classification approach.

MVPA techniques such as time-resolved decoding [10]–[13] and temporal generalization [14], have a focus on how cognitive processes unfold over time, providing evidence on the temporal evolution of the processing of specific types of stimuli and how this is classifiable as compared to others types. The neural temporal dynamics shaped by numeracy and literacy have not been explored by taking advantage of these methods. If number and letter processing are recruiting similar neural networks, we would expect chance-level classification accuracy in our study; on the other hand, if the neural dynamics for these two types of stimuli are different, their classification accuracy should be significantly higher than chance. Time-resolved decoding could provide evidence on the time intervals in which numbers and letters processing are more dissociable; temporal generalization will provide evidence on how stimulus-specific neural activity could generalize across time intervals.

The present study addressed two experimental questions. First, can the neural correlates of the visual perception of a specific class of stimuli (either numbers or letters) be successfully classified as compared to (i) visually similar but not meaningful stimuli (false fonts) and (ii) the other stimulus category (i.e., letters and number respectively)? Second, is this classification mediated by the fact that such stimuli were presented in isolation or in a string (see [4], [6])?

We used sensor-level MEG to compare the neural representations for numbers, letters, and the relative false fonts (i.e., for both numbers and letters). Participants were presented with visual stimuli (either single characters in Experiment 1 or strings of characters in Experiment 2) and required to perform a low-level visual task (pressing a button whenever a dot appeared on the screen). We selected a low-level visual task to avoid participants get involved in highly specific cognitive operations that could be qualitatively different for numbers and letters.

## Methods

### Participants

The univariate analysis of the data used in this article has been previously published in [15]. However, the multivariate analysis presented in this paper is unpublished and novel. Twenty participants (mean age: 24 ± 3years, all right-handed) were selected via the BCBL participant recruitment system (*web Participa*). All the participants had normal or corrected to normal vision and were free from any neurological or psychological brain disorder. All participants gave their written informed consent following guidelines approved by the Research Committees of the Basque center on Cognition, Brain and Language.

### Experimental procedure

All the participants took part in two experiments performing a visual detection task (i.e., white dot detection). Both experiments were presented using Psychtoolbox [16]. In the first experiment, participants were presented with Numbers (1–9), nine capital Letters (i.e., A, C, D, F, L, P, S, U, and V), and their corresponding false fonts (i.e., false-Numbers font and false-Letter font). The false fonts were created using a process similar to Shum et al. 2013 [4] keeping angles, curves, and the number of pixels as similar as possible. It was ensured that the false fonts were unrecognizable. The stimuli were presented in the center of the screen (screen placed close to ∼1 meter away from the participant’s eyes) in a white-colored Arial font with a grey background. Each stimulus (i.e., Numbers, Letters, false-Number font, false-Letter font) was repeated 22 times resulting in a total of 198 stimuli per condition (trials). Each trial started with a 500 ms baseline period, followed by a stimulus presented for 500 ms. A variable interstimulus interval (1000 – 1500 ms ISI) was introduced between trials. The participants were asked to blink during the interstimulus interval to minimize the eye blink artifacts. To keep the participants focused and attentive to the experimental manipulations, catch trials (10% of total trials) consisting of a white dot in the centre of the screen were introduced. Participants were asked to press a button. These catch trials were not included in any subsequent analyses.

In the second experiment, stimuli were created using 5-6 Letter combinations or Numbers (1– 9) separately. Phototactically legal pseudowords were used instead of consonant strings to reduce any effect of context, vocabulary, or prior probabilities in the experiment. The pseudowords were the following: BOIRA, DOCHAS, ASIMA, MODRO, DOBECA, TEPOR, PLETAR, TOLAS, EGALO.

### MEG Data Recordings

MEG data were acquired in a magnetically shielded room (to avoid environmental magnetic interference) using the whole head MEG system (MEGIN-Elekta Neuromag, Finland) installed at BCBL (https://www.bcbl.eu/en/infrastructure-equipment/meg). The MEG system consists of 102 triplet sensor pairs (each sensor consists of 102 magnetometers and 204 gradiometers arranged in a helmet configuration). The head position and movement were continuously monitored with five Head positioning coils (HPI) during the experiment. The location of HPI coils was defined relative to three fiducial points (nasion, left preauricular (LPA) and right preauricular point (RPA)) using a 3D head digitizer (Fastrack Polhemus, Colchester, VA, USA). This process is critical for correcting movement artifacts (movement compensation) during the data acquisition. The MEG data were recorded continuously with a bandpass filter (0.01–330 Hz) and a sampling rate of 1 kHz. Eye movements were monitored with pairs of electrodes placed above and below the eye (horizontal electrooculogram - HEOG) and on the external canthus of both eyes (vertical electrooculogram - VEOG).

Similarly, an electrocardiogram (EKG) was also recorded using two electrodes (in bipolar montage) placed on the right side of the abdomen and below the left clavicle of the participant. The continuous MEG data were preprocessed offline to suppress the external electromagnetic field using the temporal Signal-Space-Separation (tSSS) [17] method implemented in Maxfilter software (MaxFilter 2.1). The MEG data were also corrected for movement compensation, and the bad channels were repaired within the MaxFilter algorithms. All the analyses were carried out in Matlab version 2014B.

### Data Preprocessing

MEG data were further preprocessed using the Fieldtrip toolbox for EEG/MEG analysis [18]. The data were corrected for SQUID jump and muscle artifacts using automatic threshold-based methods implemented in the Fieldtrip toolbox. Independent components analysis (ICA) was used to remove the eye-blink (EOG) and electrocardiogram (EKG) artifacts from MEG data (using the *‘runica’* algorithm implemented in the fieldtrip). On average two components were removed for every participant, these components were visually inspected before rejection. The data were then bandpass filtered in the range of 0.5–35 Hz, demeaned, and segmented into shorter epochs ranging from 100 ms before and 500 ms after the presentation of stimuli. MEG data were then downsampled to 200 Hz. The data were baseline corrected in the range of 100 ms before the stimulus onset.

### Time-resolved decoding

Time-resolved decoding (also known as the temporal decoding approach) was used to decode between Numbers and Letters perception and false fonts. Three decoding models were generated in both experiments a) Numbers versus False-Number fonts; b) Letters versus False-Letter fonts; c) Numbers versus Letters. The data were then fed to the classifier separately for each model to perform decoding using a logistic regression classifier (LDA) implemented in the MVPA Light toolbox [19]. The classification was performed separately at each time point in both experiments. We used k-fold cross-validation (k=5) to avoid overfitting and used the area under the ROC curve (*AUC*) for evaluating the classification performance. We also used an ‘*undersampling*’ approach to balance the classes before feeding the data into a classifier for learning. This model learning process was repeated twenty (20) times for each subject to yield a stable decoding pattern. To evaluate the statistical significance of decoding across time, cluster-corrected sign permutation tests [13], [20]–[22] (i.e., one-tailed) were applied to the ‘AUC’ values obtained from the classifier with a cluster-defining threshold (i.e., alpha) p < 0.05, cluster-alpha threshold p < 0.05 and 50000 permutations considering the chance level decoding 0.50. To evaluate the statistical decoding differences across experiments, the difference in ‘AUC’ for each condition (Experiment 2 – Experiment 1) was computed. The cluster-corrected sign permutation tests [13], [20]–[22] (one-tailed) were applied to the differential ‘AUC’ values across experiments cluster-defining threshold (i.e., alpha) p < 0.05, cluster-alpha threshold p < 0.05 and 50000 permutations considering the chance level decoding 0.00.

### Temporal Generalization

Time-resolved decoding helps to understand how cognitive processes unfold over time; however, this method does not inform about the temporal organization of different stages of information processing in the human brain. To do so, the temporal generalization method is recommended, which measures the ability of a classifier to generalize across time [14]. This method provides a novel way to understand how mental representations are manipulated and transformed in a given cognitive process. In our experiment, the data pre-processing steps (i.e., low pass filtering (25 Hz) and down-sampling (200 Hz)) followed by a Logistic regression classifier with five-fold cross-validation were used as explained in the previous section. Here a classifier is trained on each time point in the data and then tested over all the available time points separately for each experimental contrast. The classifier output results in a square matrix per block per participant, where each point reflects the generalization strength of the classifier trained at a given time point and tested at another one. To evaluate the presence of a group-level effect, a cluster-based permutation test was used with cluster-defining threshold (i.e., alpha) p < 0.05, cluster-alpha threshold p < 0.05, and 10000 permutations.

## Results

### Temporal decoding

First, the time-resolved decoding was compared for conditions (i.e., Numbers versus False Numbers fonts; Letters versus False Letter fonts; and Numbers versus Letters) that were plotted separately for Experiment 1 and Experiment 2. Figure 2a shows the decoding in Experiment 1, the decoding curve for Numbers versus False Number fonts is reliably different from the chance level starting at 105 ms and achieving a peak decoding strength of 0.64 at 225 ms after the presentation of the stimuli. The decoding curve for Letters versus False Letter fonts is significantly different from the chance level starting at 110 ms and achieving a peak decoding of 0.58 at 220 ms after the presentation of the stimuli. Finally, the decoding curve between Numbers and Letters is reliable starting at 110 ms and achieving a peak accuracy of 0.55 at 160 ms after the presentation of the stimuli.

**Figure 1:**
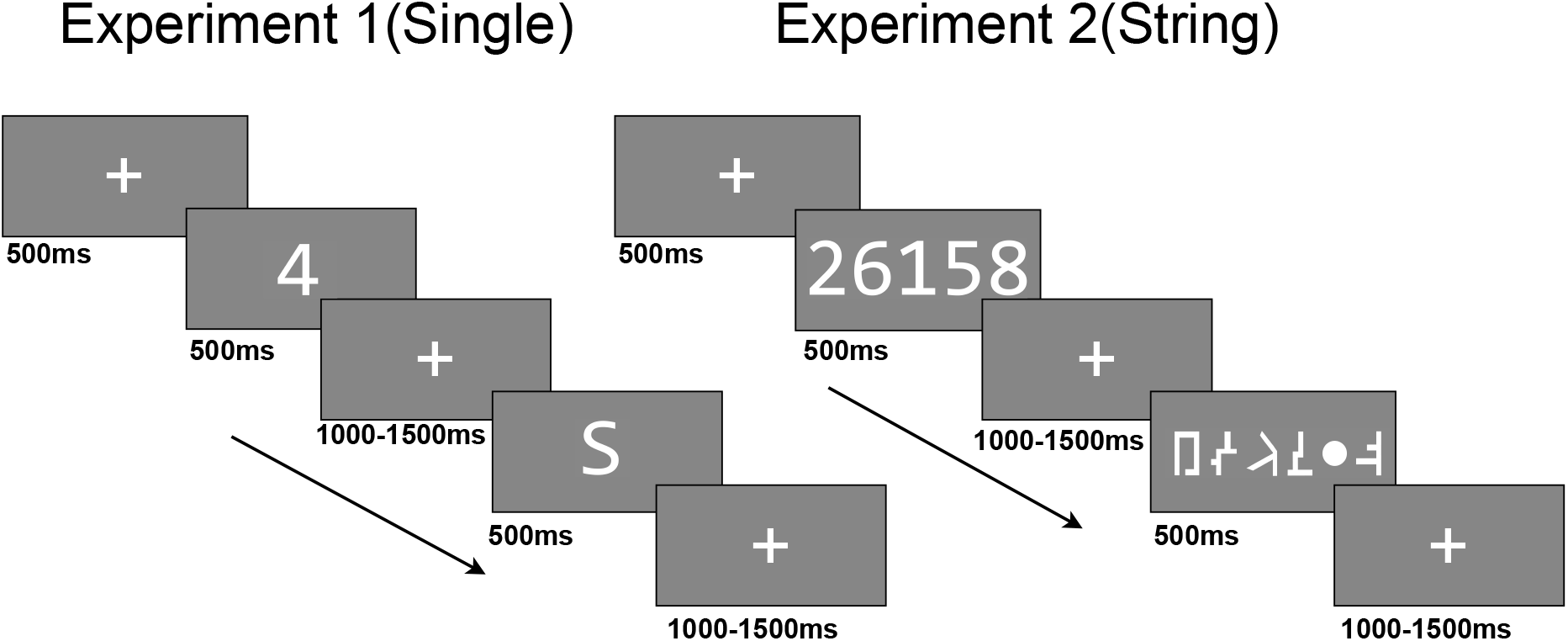
Visual detection task for single characters (Left) and sequence of characters (Right). Participants were asked to report the white dot presented randomly.

**Figure 2:**
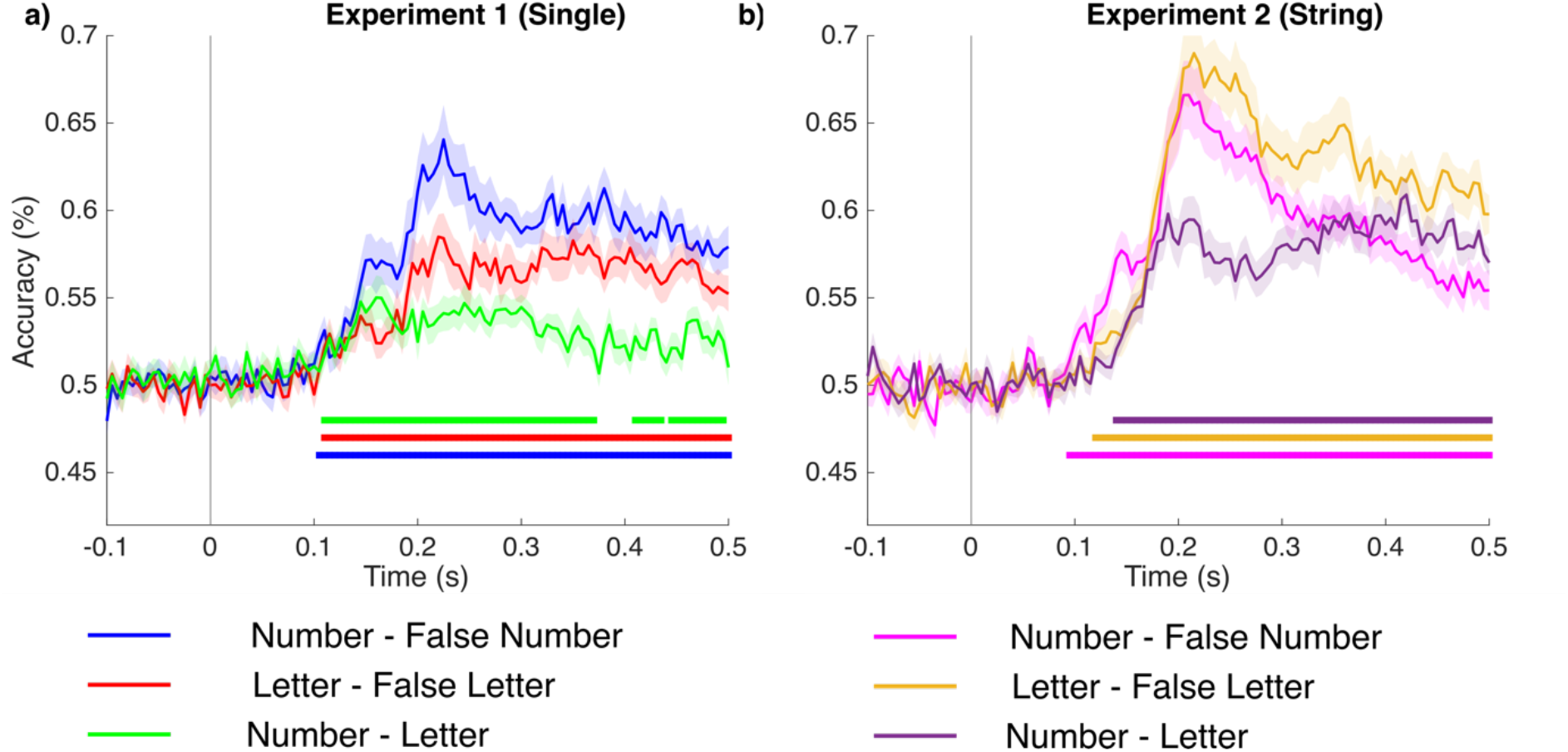
Time-resolved decoding for a) single characters for all three comparison conditions, i.e., Numbers versus False Number fonts; Letters versus False Letter fonts; and single Numbers versus single Letters (Experiment 1), and b) the same contrasts for strings of characters (Experiment 2). The colored lines show the statistically significant time points using a cluster-based permutation test (*p <* 0.05).

In Experiment 2, (Figure 2b), the decoding curve for the Number strings versus False Number font strings differs from chance starting at 95 ms and achieving a peak decoding strength of 0.66 at 210 ms after the presentation of the stimuli. The decoding curve for the Letter strings versus the False Letter font strings is reliable starting at 120 ms and achieving a peak decoding of 0.68 at 215 ms after the presentation of stimuli. Finally, the decoding curve between the Number strings and the Letter strings is significant starting at 140 ms and achieving a peak accuracy of 0.60 at 420 ms after the presentation of stimuli.

Further, to evaluate the difference between Experiment 1 (single characters) and Experiment 2 (strings), all three comparisons were plotted together. Figure 3a shows the decoding curves for Numbers versus false Numbers in Experiments 1 and 2. The grey line shows the statistically significant time points where the two curves differed statistically. The difference between the curve lasts for 40 ms (180 – 220 ms) and the difference in peak decoding strength difference is nearly 2 %. The difference between decoding curves for Letter versus false Letters in Experiments 1 and 2 emerges at 165 ms and lasts up to 500 ms with a peak decoding strength difference of almost 10 % (Figure 3c). The difference between decoding curves for Numbers versus Letters in Experiments 1 and 2 emerges at 175 ms and lasts up to 250 ms and then again emerges at 285 ms lasting up to 500 ms with a peak decoding strength difference of nearly 5% (Figure 3b).

**Figure 3:**
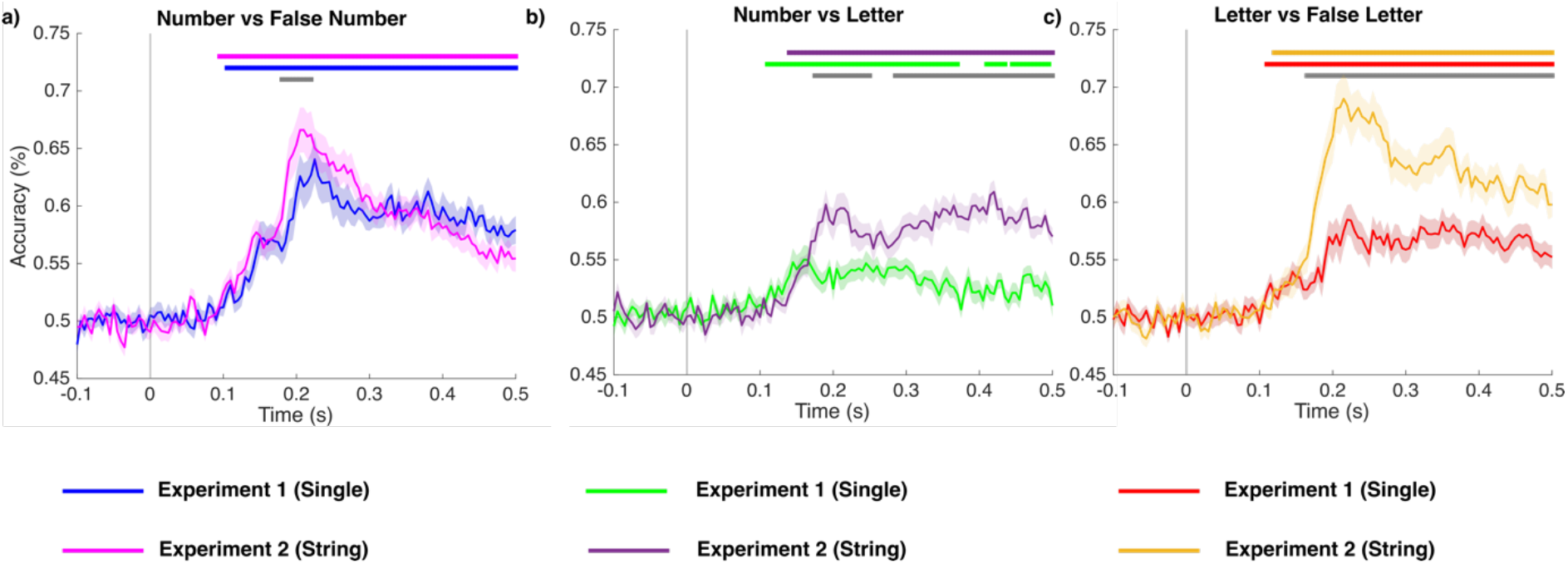
Time-resolved decoding for Numbers versus False Number fonts (Left); Letters versus False Letter fonts (Right); and Numbers versus Letters (Middle) for single characters (Experiment 1) and strings of characters (Experiment 2). The colored lines show the statistically significant time points using a cluster-based permutation test (*p <* 0.05). The grey line shows the statistically significant difference (Experience 2 – Experience 1) between single characters (Experiment 1) and a sequence of characters (Experiment 2).

### Temporal generalization

Temporal generalization matrices for Numbers versus False Numbers fonts in strings (Experiment 2) have stronger generalization across diagonal compared to single Numbers versus false Number fonts (Experiment 1). The generalization for single characters starts at 90 ms and becomes strongest between 200 and 300 ms and sustains up to 500 ms. For the strings (Experiment 2), it generalizes starting from 90 ms and peaks between 200 and 400 ms. The difference between generalization matrices (Experiment 2 – Experiment 1) does not show statistical differences across the diagonal (upper row of matrices in Figure 4). Interestingly though, the decoding classifier trained at around 200 ms can generalize and successfully classify data from 200 to 400 ms.

**Figure 4:**
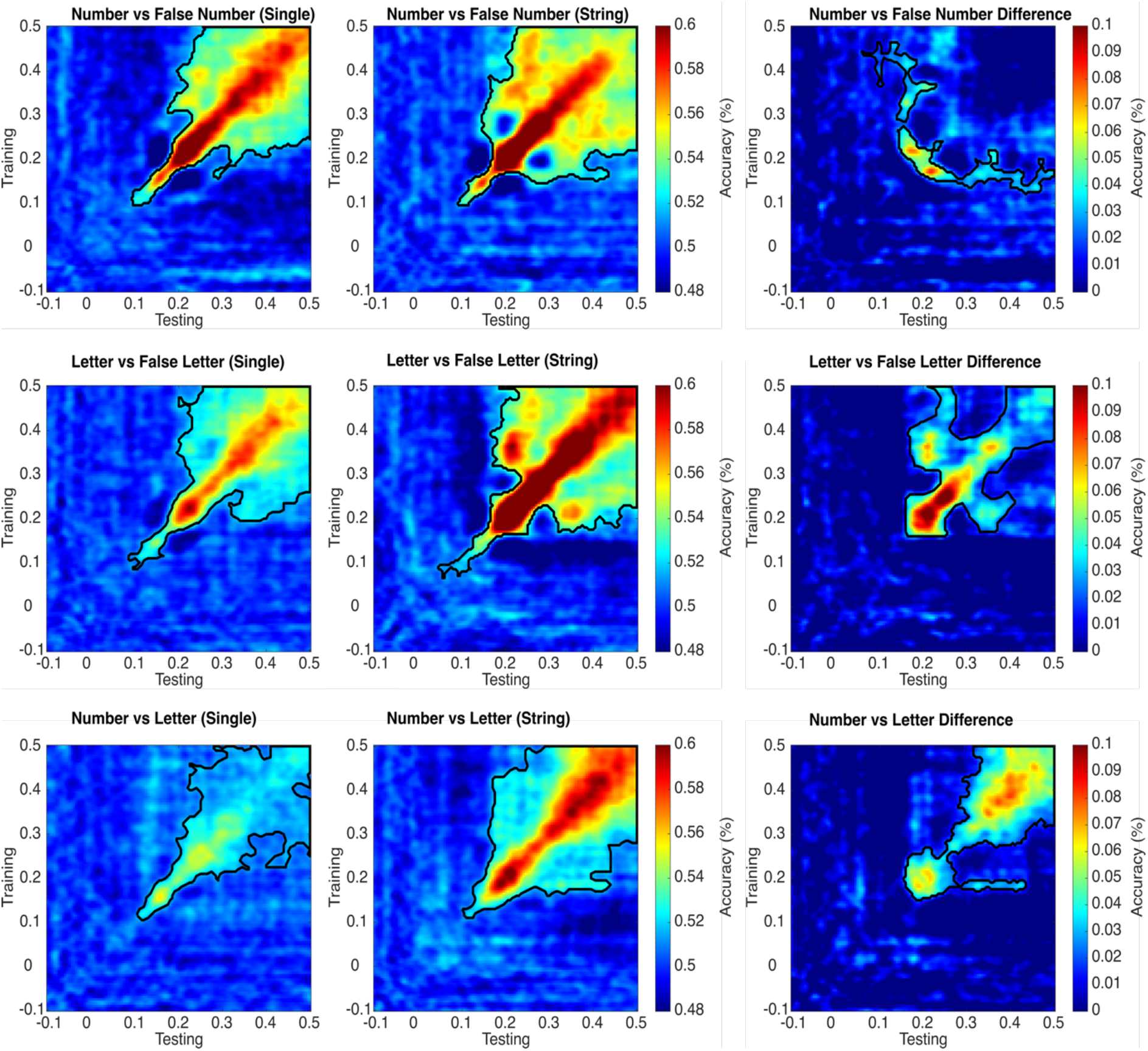
Temporal generalization plots for Numbers versus False Number font in single characters i.e., Experiment 1 (first row first column); Letters versus False Letter fonts (second row first column); and Numbers versus Letters (third row first column). The second column represents similar comparisons to first column but for the string of characters (Experiment 2). The third column represents the difference between single and string of characters (Experiment 2 – Experiment 1). The black colored outline shows the boundaries of statistically significant time points using a cluster-based permutation test (*p <* 0.05).

For single (Experiment 1) Letters versus False Letter fonts, the generalization starts from 85 ms and peaks between 200 and 300 ms, whereas for the strings (Experiment 2), generalization starts as early as 45 ms and peaks from 200 to 500 ms. The diagonal generalization plot is stronger in the strings compared to the single characters. The generalization difference (Experiment 2 – Experiment 1) starts at 160 ms, peaks between 200 and 300 ms, and appears to be sustained up to 500 ms (middle row of matrices in Figure 4).

For single (Experiment 1) Numbers versus Letters, the generalization starts from 105 ms and peaks between 200 and 300 ms. This generalization is weaker compared to the other two comparisons i.e., Numbers versus False Numbers and Letters versus False Numbers. For strings (Experiment 2), the generalization starts from 100 ms and first peaks between 150 and 250 ms and then again peaks between 300 and 500 ms. The generalization difference (Experiment 2 – Experiment 1) starts at 155 ms and peaks between 300–500 ms. Also, the classifier trained close to 200 ms can generalize when testing from 250 to 450 ms (lower row of matrices in Figure 4).

## Discussion

In these two experiments, we report logistically dissociable temporal dynamics for processing numbers and letters in the human brain. We observed a significant difference in decoding accuracy for both numbers and letters compared to the respective false fonts and between each other directly. We also observed that the decoding accuracy was highly mediated by the fact whether the stimuli were presented in isolation (i.e., single characters, Experiment 1) or as a string (i.e., a string of characters, Experiment 2) and this was especially true for letters.

Concerning our first experimental question (are numbers and letters functionally dissociable?), our results indicate that numbers and letters are processed differently both when presented as single/isolated and when in a string. When presented in isolation, the neural representation of numbers (vs. visually similar false fonts) is stronger (i.e., triggered higher decoding accuracy) if compared to the decoding accuracy for letters (vs. false letter fonts). When the stimuli are a string of similar characters, however, the decoding accuracy for letters is stronger in magnitude compared to the numbers (vs. false fonts). For both single items and strings, the direct classification of numbers vs. letters processing showed lower decoding accuracy, as compared to the individual comparisons with false fonts (Figure 2). The temporal generalization results tell a similar story. Numbers and letters representations were stronger when compared to their respective false fonts but when compared across different categories (i.e., number versus letter), the generalization patterns were slightly weaker. It thus appears that one important component triggering such neural dissociation (letters or numbers vs. their respective false fonts), even in such an early time window (∼100 ms), is the culturally determined visual familiarity with the visual input. When directly comparing the two types of stimuli (letters vs. numbers), the neural dissociation is weaker, probably because a relevant portion of the neural processes at work for those familiar stimuli is shared.

Importantly, however, our results highlight that the number and letter processing is highly affected by whether these characters were presented either in isolation as a string (Figure 3). Number processing is not very affected by the presentation mode (either single or strings). The difference between the two presentation modes was only present around the peak decoding and was also short-lived (∼40 ms, Figure 3a). On the other hand, letters were strongly affected by the presentation mode. Letters presented as strings of characters were highly decodable (∼20 % higher peak decoding accuracy, compared to single letters): this difference across experiments started from 165 ms and was sustained for the whole window of interest (Figure 3c). Along these lines, it is worth remarking that decoding accuracy for numbers was higher when stimuli were presented in isolation (Figure 2a), while decoding for letters was higher when stimuli were strings (Figure 2b). This is mainly due to the large across-experiment decoding difference observed for letters. Finally, numbers vs. letters decoding was stronger in experiment 2 compared to experiment 1 (Figure 3b), even if decoding accuracy for the two classes of items compared to the respective false fonts was more similar in this second experiment (Figure 2b).

The temporal generalization plots (in Figure 4) confirm the pattern of results discussed for temporal decoding. Here however, we can appreciate i) the generalization patterns across different time points and consequently ii) if some neural mechanisms emerging across time can be functionally dissociated. Concerning the first point, the generalization plots for strings of letters indicate the presence of a recurrent neural process that can generalize from the activity ∼200 ms in the diagonal to later activity ∼350 ms, an effect that is not evident for single letters (second row in Figure 4). Such an effect is not present for numbers (first row in Figure 4). Concerning the second point, when comparing numbers and letters directly (third row in Figure 4), there is evidence for two time intervals (early ∼200 ms and later ∼400 ms across the diagonal) reflecting independent processing steps, that are present for strings and not for single items.

Based on these findings, it emerges that the combinatorial mechanisms for numbers and letters are key for triggering dissociable neural patterns for the two categories of stimuli, at least when participants are involved in a low-level perceptual task. The presentation of numbers either in isolation or in a string does not change drastically the quality of neural processes at work. On the other hand, letters in isolation elicit different neural processes compared to letter strings (pseudowords in our experiment). In other words, seeing isolated alphabetic stimuli, such as “*a*”, “*c*” and “*t*”, does not trigger the same cognitive processes compared to a word such as “*cat*”, or a pseudoword such as “*tac*” (as in Experiment 2). Such stimuli elicit early letter-specific neural activity (∼200 ms) that is then reactivated later in time (∼350 ms), probably feeding hierarchically more abstract neurocognitive processes at ∼400 ms, that have been classically associated with language specific lexical/semantic analysis of the printed stimuli. In fact, phonotactically legal pseudowords can elicit the activation of a larger cohort of orthographical and phonological similar words (neighbors) in the human mental lexicon, as compared to real words [23]. Park and colleagues [6] observe “*that the visual cortex is tuned to selectively process combinations of letters, but not numbers, further along in the visual processing stream*”. The present results are in line with this observation. It is possible that stimulating more abstract mental operations (with specific arithmetic tasks) would trigger later neural activity also for numbers and that such activity could be dissociable for single compared to strings of numbers. Even so, it is important to observe that such activity is not automatically elicited in our low-level detection task, while this is the case for strings of letters. Thus, the present data speak for a clear neural dissociation between numeracy and literacy, mainly when presenting the stimuli in strings, but not so much when numbers and letters are presented in isolation.

Previous studies [4], [8], [15] mainly focused on the lateralization effect between number and letter processing. Park and colleagues [6] investigated the time course of the dissociation between visually-presented letters and numbers and reported highly similar temporal effects to the ones we observed in the present analysis. A hemispheric dissociation between the ERP waveforms evoked by numbers and letters at the occipital-temporal electrodes was observed. Specifically, letters elicited significantly greater N1 amplitudes (∼150 ms) in the left hemisphere, while numbers elicited significantly greater N1 amplitudes in the right hemisphere (especially for strings). However, this evidence of a different hemispheric sensitivity for the two types of stimuli does not preclude that both hemispheres are involved in the processing of numbers and letters. One potential explanation for the hemispheric dissociation observed by Park and colleagues (2014) could be the fact that one of the two categories of stimuli is more complex to process, hence triggering higher neurocognitive demands for the same neural mechanisms at work for the two categories of stimuli. Research by Shum and colleagues [4] and by Carreiras and colleagues [8] showed a right hemisphere involvement for numbers, but no clear dissociation in the recruitment of the left hemisphere. Univariate analysis of the present data [15] did not provide clear support for the hemispheric dissociation either. The classical univariate analysis would not be suited for dissociating different sources of variability since the individual brain responses are averaged across multiple trials and the fine-grained trial-by-trial variability would be lost. We used a machine learning approach ([14], [24]) to explore the trial-by-trial variability of the neural responses to numbers and letters and thus evaluated the stronger hypothesis that the neural responses to numbers and letters can be logistically classified into separate categories. This analysis revealed, across time points, that the functional response to numbers and letters can be accurately decoded mainly for strings, thus showing when the neural activity becomes number-or letter-specific. Given the early decoding for numbers vs. letters observed in the present study (starting ∼100 ms), we can thus conclude that the presentation of number and letter strings recruits dissociable perceptual neural networks in the human brain. Our results do not focus on hemispheric lateralization, but based on previous evidence [4], [15] we can advance that the two hemispheres (and especially the left one) are recruited for both letters and numbers, with more fine-grained functional dissociations in the micro-neural networks recruited by the two categories of stimuli.

The present study thus provides interesting experimental evidence on the effects of literacy vs. numeracy that can stimulate further research. First, we have reported a significant difference in decoding patterns for letters while presented in strings compared to strings of numbers. Second, we reinforce the idea that early neural activity of the human cortex is fine-tuned to selectively process combinations of letters (as compared to single letters) along the visual processing stream. Third, we reported such dissociations in a low-level visual detection task, where no differential task demands for numbers of letters can explain the observed dissociations. Further research, maybe involving intracranial recordings, should further explore the differential role of the visual pathways in the two hemispheres, thus clarifying the spatial dissociation supporting the functional dissociation we report.

## Acknowledgments

This research was supported by the Basque Government through the BERC 2018–2021 program and by the Spanish State Research Agency through BCBL’s Severo Ochoa excellence accreditation CEX2020-001010-S and the project BES-2016-077560 funded by the Spanish Ministry of Economy and Competitiveness (MINECO). SN acknowledges the support from “The Adaptive Mind,” funded by the Excellence Program of the Hessian Ministry of Higher Education, Science, Research and Art. MC was supported by “la Caixa” Foundation (ID 100010434), under the agreement HR18-00178-DYSTHAL, and by the Agencia Estatal de Investigación PID2021-122918OB-I00. NM was supported by the Spanish Ministry of Science, Innovation and University (grants PSI2015-65694-P, RTI2018-096311-B-I00), the Agencia Estatal de Investigación (AEI), the Fondo Europeo de Desarrollo Regional (FEDER) and by the Basque government (grant PI_2016_1_0014). HR was supported by the Economic and Social Research Council (ESRC) funded Business and Local Government Data Research Centre under Grant ES/S007156/1.

SN thanks the BCBL lab research staff for their valuable support.

## Author contributions

**S.N.**: Conceptualization, Methodology, Software, Data curation, Writing – original draft. **H.R**.: Methodology, Visualization, Writing – review & editing, Funding acquisition. **MC**: Methodology, Supervision, Writing – review & editing. **NM**: Conceptualization, Methodology, Validation, Supervision, Funding acquisition, Project administration, Writing – review & editing.

## Data availability statement

The raw data can be accessed from Open-Neuro repository (https://openneuro.org/datasets/ds002712/versions/1.0.1). Due to the whole preprocessed dataset’s size and limited public storage options available, only raw data has been uploaded. However, the full preprocessed dataset is available upon requests directed to Dr. Nicola Molinaro or Dr. Sanjeev Nara. The full data set could then be shared through the private BCBL-secured institutional servers temporarily available for big data transfer.

## References

[1] S. Dehaene et al., ‘How learning to read changes the cortical networks for vision and language’, Science, vol. 330, no. 6009, pp. 1359–1364, Dec. 2010, doi: 10.1126/SCIENCE.1194140.

[2] S. Dehaene, L. Cohen, J. Morais, and R. Kolinsky, ‘Illiterate to literate: behavioural and cerebral changes induced by reading acquisition’, Nature Publishing Group, vol. 16, 2015, doi: 10.1038/nrn3924.

[3] S. Caffarra, M. Lizarazu, N. Molinaro, and M. Carreiras, ‘Reading-Related Brain Changes in Audiovisual Processing: Cross-Sectional and Longitudinal MEG Evidence’, Journal of Neuroscience, vol. 41, no. 27, pp. 5867–5875, Jul. 2021, doi: 10.1523/JNEUROSCI.3021-20.2021.

[4] J. Shum et al., ‘A Brain Area for Visual Numerals’, 2013, doi: 10.1523/JNEUROSCI.4558-12.2013.

[5] J. Park, A. Hebrank, T. A. Polk, and D. C. Park, ‘Neural Dissociation of Number from Letter Recognition and Its Relationship to Parietal Numerical Processing’, J Cogn Neurosci, vol. 24, no. 1, p. 39, Jan. 2012, doi: 10.1162/JOCN_A_00085.

[6] J. Park, C. Chiang, E. M. Brannon, and M. G. Woldorff, ‘Experience-Dependent Hemispheric Specialization of Letters and Numbers is Revealed in Early Visual Processing’, J Cogn Neurosci, vol. 26, no. 10, p. 2239, Oct. 2014, doi: 10.1162/JOCN_A_00621.

[7] J. Park, ‘A neural basis for the visual sense of number and its development: A steady-state visual evoked potential study in children and adults’, Dev Cogn Neurosci, vol. 30, pp. 333–343, Apr. 2018, doi: 10.1016/J.DCN.2017.02.011.

[8] M. Carreiras, P. J. Monahan, M. Lizarazu, J. A. Duñabeitia, and N. Molinaro, ‘Numbers are not like words: Different pathways for literacy and numeracy’, 2015, doi: 10.1016/j.neuroimage.2015.06.021.

[9] M. Grotheer, K. H. Herrmann, and G. Kovács, ‘Neuroimaging Evidence of a Bilateral Representation for Visually Presented Numbers’, Journal of Neuroscience, vol. 36, no. 1, pp. 88–97, Jan. 2016, doi: 10.1523/JNEUROSCI.2129-15.2016.

[10] R. M. Cichy, D. Pantazis, and A. Oliva, ‘Resolving human object recognition in space and time’, Nat Neurosci, vol. 17, no. 3, pp. 455–462, Mar. 2014, doi: 10.1038/nn.3635.

[11] S. Nara et al., ‘Temporal uncertainty enhances suppression of neural responses to predictable visual stimuli’, Neuroimage, vol. 239, Oct. 2021, doi: 10.1016/j.neuroimage.2021.118314.

[12] T. Grootswagers, S. G. Wardle, and T. A. Carlson, ‘Decoding dynamic brain patterns from evoked responses: A tutorial on multivariate pattern analysis applied to time series neuroimaging data’, J Cogn Neurosci, vol. 29, no. 4, pp. 677–697, Apr. 2017, doi: 10.1162/jocn_a_01068.

[13] S. Nara, D. Rathee, N. Molinaro, N. du Bois, B. Bhushan, and G. Prasad, ‘Temporal Dynamics of Neural Processing of Facial Expressions and Emotions’, bioRxiv, p. 2021.05.12.443280, Jan. 2021, doi: 10.1101/2021.05.12.443280.

[14] J. R. King and S. Dehaene, ‘Characterizing the dynamics of mental representations: The temporal generalization method’, Trends in Cognitive Sciences, vol. 18, no. 4. Elsevier Ltd, pp. 203–210, 2014. doi: 10.1016/j.tics.2014.01.002.

[15] S. Aurtenetxe, N. Molinaro, D. Davidson, and M. Carreiras, ‘Early dissociation of numbers and letters in the human brain’, Cortex, vol. 130, pp. 192–202, Sep. 2020, doi: 10.1016/j.cortex.2020.03.030.

[16] D. H. Brainard, ‘The Psychophysics Toolbox’, Spat Vis, vol. 10, no. 4, pp. 433–436, 1997, doi: 10.1163/156856897X00357.

[17] S. Taulu and J. Simola, ‘Spatiotemporal signal space separation method for rejecting nearby interference in MEG measurements’, Phys Med Biol, vol. 51, no. 7, pp. 1759– 1768, Apr. 2006, doi: 10.1088/0031-9155/51/7/008.

[18] R. Oostenveld, P. Fries, E. Maris, and J. M. Schoffelen, ‘FieldTrip: Open source software for advanced analysis of MEG, EEG, and invasive electrophysiological data’, Comput Intell Neurosci, vol. 2011, 2011, doi: 10.1155/2011/156869.

[19] M. S. Treder, ‘MVPA-Light: A Classification and Regression Toolbox for Multi-Dimensional Data’, Front Neurosci, vol. 14, Jun. 2020, doi: 10.3389/fnins.2020.00289.

[20] E. Maris and R. Oostenveld, ‘Nonparametric statistical testing of EEG- and MEG-data’, J Neurosci Methods, vol. 164, no. 1, pp. 177–190, Aug. 2007, doi: 10.1016/j.jneumeth.2007.03.024.

[21] S. Nara, M. Lizarazu, C. G. Richter, D. C. Dima, M. Bourguignon, and N. Molinaro, ‘Temporal uncertainty affects the visual processing of predicted stimulus properties’.

[22] D. C. Dima and K. D. Singh, ‘Dynamic representations of faces in the human ventral visual stream link visual features to behaviour’, 2018.

[23] M. Kutas and K. D. Federmeier, ‘Thirty years and counting: Finding meaning in the N400 component of the event related brain potential (ERP)’, Annu Rev Psychol, vol. 62, p. 621, Jan. 2011, doi: 10.1146/ANNUREV.PSYCH.093008.131123.

[24] J. R. King, A. Gramfort, A. Schurger, L. Naccache, and S. Dehaene, ‘Two distinct dynamic modes subtend the detection of unexpected sounds’, PLoS One, vol. 9, no. 1, Jan. 2014, doi: 10.1371/journal.pone.0085791.

